# Association of mitochondria with microtubules inhibits mitochondrial fission by precluding Dnm1 assembly

**DOI:** 10.1101/178913

**Authors:** Kritika Mehta, Leeba Ann Chacko, Manjyot Kaur Chug, Siddharth Jhunjhunwala, Vaishnavi Ananthanarayanan

## Abstract

Mitochondria are organized as tubular networks in the cell and undergo fission and fusion. While several of the molecular players involved in mediating mitochondrial dynamics have been identified, the precise cellular cues that initiate fission or fusion remain largely unknown. In fission yeast, mitochondria are organized along microtubule bundles. Here, we employed deletions of kinesin-like proteins to perturb microtubule dynamics, and determined that cells with long microtubules exhibited long, but fewer mitochondria, whereas cells with short microtubules exhibited short, but several mitochondria due to reduced mitochondrial fission in the former and elevated fission in the latter. Correspondingly, upon onset of closed mitosis in fission yeast, wherein interphase microtubules assemble to form the spindle within the nucleus, we measured increased mitochondrial fission. We determined that the consequent rise in the mitochondrial copy number was necessary to reduce partitioning errors while stochastically partitioning mitochondria between daughter cells. We discovered that the association of mitochondria with microtubules physically impeded the assembly of the fission protein Dnm1 around mitochondria, resulting in inhibition of mitochondrial fission. Taken together, we demonstrate a novel mechanism for regulation of mitochondrial fission that is dictated by the interaction between mitochondria and the microtubule cytoskeleton.

## Introduction

Mitochondria are double-membraned organelles whose functions range from ATP production to calcium signalling. Inside cells, mitochondrial form is dynamic and transitions from tubular networks to fragmented entities depending on the activity of the mitochondrial fission and fusion machinery. The major mitochondrial fission protein is dynamin-related Drp1 GTPase (Smirnova et al., 2001) (Dnm1 in yeast (Bleazard et al., 1999; Jourdain et al., 2009)). Multimeric Drp1 rings assemble around the mitochondrial membrane and utilize the energy from GTP hydrolysis to catalyze the constriction and fission of mitochondria (Basu et al., 2017; Ingerman et al., 2005). Fusion of mitochondria requires two sets of proteins namely Opa1 (Delettre et al., 2000) for the inner membrane (Mgm1 in yeast (Sesaki et al., 2003)) and Mfn1/2 (Eura et al., 2003) for the outer membrane (Fzo1 in yeast (Hermann et al., 1998)).

The requirement for dynamic mitochondria has been attributed to two primary reasons, namely quality control and energy production (Mishra and Chan, 2016). Larger/longer mitochondria resulting from fusion are hypothesized to be capable of producing more energy whereas shorter/smaller mitochondria formed following a fission event are likely to undergo mitophagy (Chen and Chan, 2009). In the case of the latter, fission could serve as an efficient mechanism to segregate and eliminate damaged mitochondria. Dysfunction of fission and fusion processes has been implicated in neurodegeneration (Deng et al., 2008; Hirai et al., 2001), cancer (Graves et al., 2012) and cardiomyopathies (Ashrafian et al., 2010), amongst a host of metabolic disorders.

In mammalian cells, mitochondria are transported along microtubule tracks by the activity of motor proteins kinesin-1 and dynein (Pilling et al., 2006). Kinesin-1 and dynein bind to the outer membrane of mitochondria via the Miro/Milton complex (Glater et al., 2006; Stowers et al., 2002; van Spronsen et al., 2013) and move mitochondria in the anterograde and retrograde direction respectively. In neuronal cells, increase in calcium levels results in the attachment of kinesin-1 motor to mitochondria via syntaphilin, which inhibits the ATPase activity of kinesin and hence leads to stationary mitochondria on neuronal microtubules (Chen and Sheng, 2013). About 70% of mitochondria in neuronal cells have been visualized in this stationary state (Kang et al., 2008). In contrast to mammalian cells, mitochondria in fission yeast do not undergo motor-driven movement along microtubules (Chiron et al., 2008; Yaffe et al., 2003). However, the protein Mmb1 has been identified to associate mitochondria with dynamic microtubules (Fu et al., 2011). Upon microtubule depolymerization using methyl benzimidazol-2-yl-carbamate (MBC), mitochondria have been observed to undergo fragmentation (Fu et al., 2011; Jourdain et al., 2009; Li et al., 2015). Additionally, mitochondrial dynamics and partitioning in fission yeast have been observed to be actin/myosin-independent processes (Jourdain et al., 2009), contrary to the mechanism of mitochondrial partitioning in budding yeast (Fehrenbacher et al., 2004).

Cells employ several strategies to reduce partitioning error of organelles during mitosis, such as ordered segregation mediated by spindle poles, or increasing copy numbers of organelles prior to cell division (Huh and Paulsson, 2011). In the latter, homogeneous distribution of the multiple organelle copies serves to increase the probability of equal partitioning stochastically. Mitochondrial inheritance has been observed to be microtubule-dependent in mammalian cells (Lawrence and Mandato, 2013). In fission yeast, mitochondrial partitioning during cell division has been proposed to be mediated by attachment of mitochondria with spindle poles (Jajoo et al., 2016; Krüger and Tolic, 2008; Yaffe et al., 2003), similar to the segregation of endosomes, lysosomes and Golgi bodies in mammalian cells (Bergeland et al., 2001; Shima et al., 1998). However, only a portion of the observed mitochondria associated with the spindle poles (Yaffe et al., 2003). Additionally, increased mitochondrial fragmentation upon the onset of mitosis has also been observed (Jourdain et al., 2009), perhaps indicating a stochastic mechanism for mitochondrial partitioning.

While the molecular players that effect fission and fusion have been identified in several systems, the cellular signals that regulate these events are largely elusive. Here, we demonstrate that mitochondria piggyback on dynamic microtubules to selectively undergo fission when microtubules depolymerize. Reorganization of interphase microtubules into the nucleus when cells prepare for division also provided the cue for increased mitochondrial fission. We quantified the number of mitochondria in mother cells immediately after formation of mitotic spindle within the nucleus and the number of mitochondria in the resulting daughter cells, and confirmed that the partitioning was indeed a good fit to a binomial distribution, indicating that the mitochondria were stochastically segregated into the two daughter cells. We determined that the presence of long and stabilized microtubules was inhibitory to unopposed fission even when Dnm1 was overexpressed. Finally, we discovered microtubule-bound mitochondria were unlikely to undergo fission due to the unavailability of space between microtubules and mitochondria for the formation of the Dnm1 ring.

## Materials and Methods

### Strains and media

The fission yeast strains used in this study are listed in Table S1. All the strains were grown in YE (yeast extract) media or Edinburg Minimal media (EMM) (Forsburg and Rhind, 2006) with appropriate supplements at a temperature of 30°C. Cells that were transformed with plasmid pREP41-Dnm1 or pREP41-Dnm1-Cterm-GFP (see Table S1) were cultured in EMM with appropriate supplements and 0.05μM thiamine for partial induction of the nmt1 promoter. Strains VA076 and VA084 were constructed by crossing PT2244 with FY20823, and Dnm1Δ with G5B respectively (see Table S1), following the random spore analysis protocol (Forsburg and Rhind, 2006).

### Plasmid transformation

Transformation of strains was carried out using the improved protocol for rapid transformation of fission yeast (Kanter-Smoler et al., 1994). In brief, cells were grown overnight to log phase in low glucose EMM, pelleted and washed with distilled water. The cells were then washed in a solution of lithium acetate/EDTA (100mM LiAc, 1mM EDTA, pH 4.9) and re-suspended in the same solution. 1μg of plasmid DNA was added to the suspension, followed by addition of lithium acetate/PEG (40% w/v PEG, 100mM LiAc, 1mM EDTA, pH 4.9) and then incubated at 30°C for 30 min in a shaking incubator. This was followed by a heat shock of 15min at 42°C. Thereafter, cells were pelleted down, re-suspended in TE solution (10mM Tris-HCl, 1mM EDTA, pH 7.5) and plated onto selective EMM plates.

### Preparation of yeast for imaging

For imaging mitochondria, fission yeast cells were grown overnight in a shaking incubator at 30°C, washed once with distilled water, and stained with 200nM Mitotracker Orange CMTMRos (ThermoFisher Scientific, Cat. #M7510) dissolved in EMM for 20min. Following this, cells were washed thrice with EMM and then allowed to adhere on lectin-coated (Sigma-Aldrich, St. Louis, MO, Cat. #L2380) 35mm confocal dishes (SPL, Cat. #100350) for 20min. Unattached cells were then removed by washing with EMM. In experiments where mitochondria were not imaged, staining with Mitotracker was omitted.

### Microtubule depolymerization

For depolymerization of microtubules, cells were treated with methyl benzimidazol-2-yl-carbamate (MBC, Carbendazim 97%, Sigma Aldrich). A stock solution with a concentration of 25mg/ml was prepared in DMSO and later diluted to a working concentration of 25μg/ml in EMM.

### Microscopy

Confocal microscopy was carried out using the InCell Analyzer-6000 (GE Healthcare, Buckinghamshire, UK) with 60×/0.7 N.A. objective fitted with an sCMOS 5.5MP camera having an x-y pixel separation of 108nm. For GFP and Mitotracker Orange imaging, 488 and 561nm laser lines and bandpass emission filters 525/20 and 605/52nm respectively were employed. Time-lapses for visualization of mitochondrial dynamics were captured by obtaining 5 Z-stacks with a 0.5μm-step size every 12s. Deconvolution was performed in images obtained using a Deltavision RT microscope (Applied Precision) with a 100×, oil-immersion 1.4 N.A. objective (Olympus, Japan). Excitation of fluorophores was achieved using InsightSSI (Applied Precision) and corresponding filter selection for excitation and emission of GFP and Mitotracker Orange. Z-stacks with 0.3μm-step sizes encompassing the entire cell were captured using a CoolSnapHQ camera (Photometrics), with 2×2 binning. The system was controlled using softWoRx 3.5.1 software (Applied Precision) and the deconvolved images obtained using the built-in setting for each channel.

### 3D visualization of deconvolved images

3D views of the microtubules and mitochondria in Movies S2, S3, S4, S8, S9, S10, S12 and S13 were obtained from deconvolved images captured in the Deltavision microscope using Fiji’s ‘3D project’ function, with the brightest point projection method and 360° total rotation with 10° rotation angle increment.

### Estimation of volume of mitochondria

Mitochondrial volume was estimated in Fiji by integrating the areas of mitochondria in thresholded 3D stacks of cells in fluorescent deconvolved images obtained using the Deltavision RT microscope. The total volume was then normalized to the mean total mitochondrial volume of wild-type cells. Individual mitochondrial volumes were estimated in the same fashion.

### Analysis of mitochondrial dynamics

Individual mitochondria were identified in each frame of the time-lapse obtained in confocal mode of the GE InCell Analyzer after projecting the maximum intensity of the 3D stack encompassing the cell, followed by mean filtering and visualization in Fiji’s ‘mpl-magma’ lookup table. Following identification of mitochondria, the ‘Measure’ function of Fiji was employed to obtain the circularity, aspect ratio and parameters of fitted ellipse. The length of the major axis of the ellipse fitted to a mitochondrion was defined as the size of that mitochondrion. The size, circularity and aspect ratio were estimated for mitochondria at each frame and each time point. Fission and fusion frequencies of mitochondria were estimated by counting the number of mitochondria identified during each frame of the time-lapse. The difference in number of mitochondria from one frame to the next was counted, with an increase being counted as fission event and decrease as fusion event. The total number of fission events and fusion events per cell were estimated and divided by the total duration of the time-lapse to obtain the fission and fusion frequencies.

### Test for fit of mitochondrial partitioning during mitosis to binomial distribution

To test the fit of mitochondrial segregation during mitosis to a binomial distribution, the data were z-transformed as previously described (Hennis and Birky, 1984). Briefly, given *n* mitochondria in the mother cell just prior to cell division, *x* and *n* − *x* mitochondria in the resulting daughter cells, *z* was given by 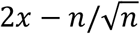 to approximate the binomial distribution to a normal distribution of 0,1. The *z* values obtained for each *x* and *n* were binned into *k* bins of equal sizes and subjected to Chi-square test with *k* − 1 degrees of freedom. The *z* values are expected to be equally distributed among the *k* bins, with expected number of 1/*k* per bin.

### Data analysis and plotting

Data analysis was performed in Matlab (Mathworks, Natick, MA). Box plots with the central line indicating the median and notches that represent the 95% confidence interval of the median were obtained by performing one-way ANOVA (‘anova1’ in Matlab) or Kruskal-Wallis Test (‘kruskalwallis’ in Matlab). The former was used when data were found to be normally distributed and the latter when data were non-normally distributed (tested using ‘chi2gof’ in Matlab). Following this, significant difference (p<0.05) was tested using the Tukey’s Honestly Significant Difference procedure (‘multcompare’ in Matlab). All the plots were generated using Matlab.

## Results

### 1. Perturbation of microtubule dynamics leads to changes in mitochondrial numbers

**Figure 1.**
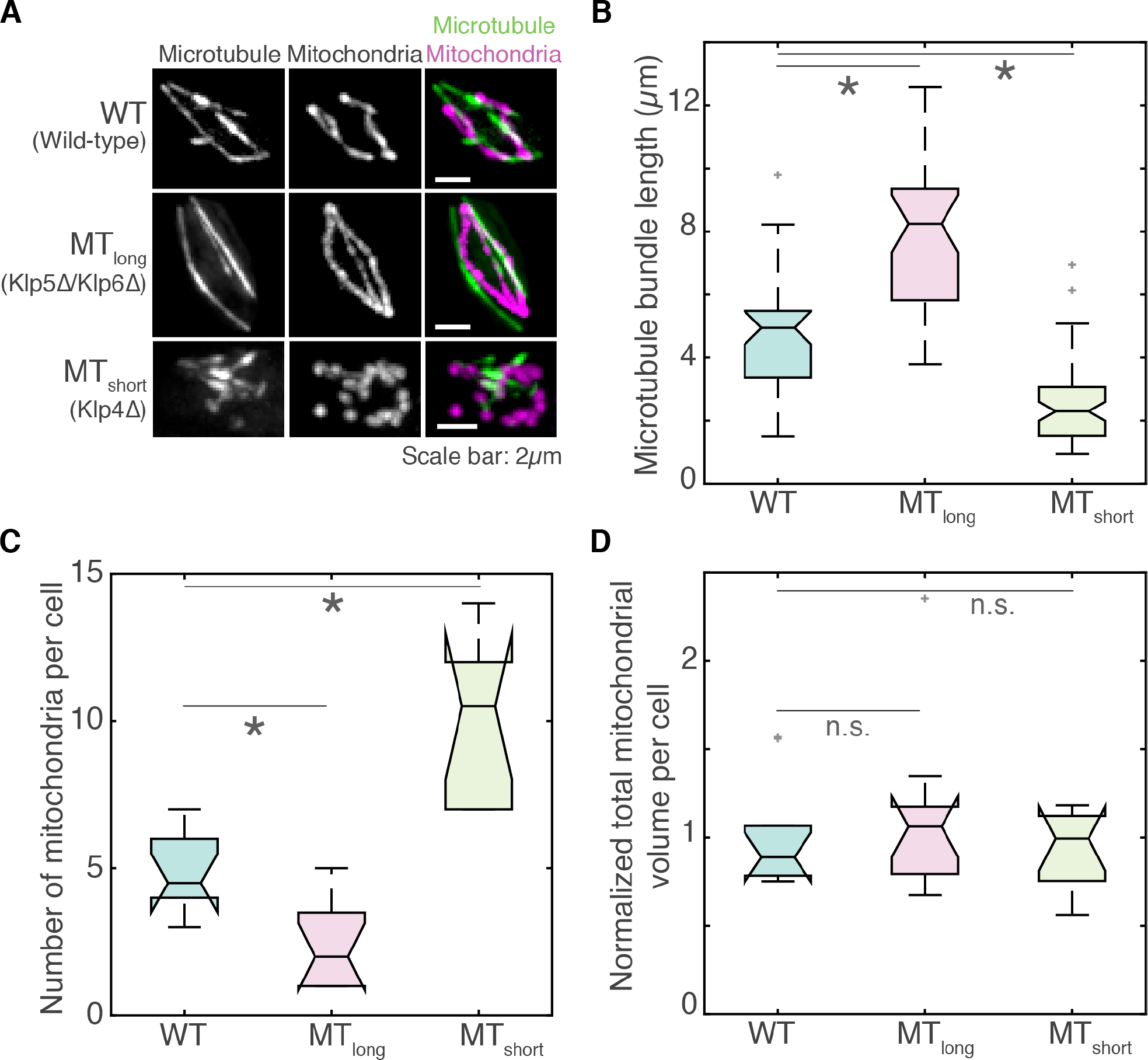
Mitochondrial number is inversely proportional to microtubule length. **(A)**Maximum intensity projections of deconvolved Z-stack images of microtubules (left), mitochondria (centre) and their composite (right) in wild-type (‘WT’, top, strain KI001, see Table S1), Klp5Δ/Klp6Δ (‘MT_long_’, strain G3B, see Table S1) and Klp4Δ (‘MT_short_’, strain G5B, see Table S1) cells. **(B)**Box plot of length of anti-parallel microtubule bundles in WT, MT_long_ and MT_short_ cells (*n*=40, 37 and 63 bundles respectively). **(C)**Box plot of number of mitochondria per cell in WT, MT_long_ and MT_short_ cells (*n*=10, 12 and 13 cells respectively). **(D)**Box plot of the total volume of mitochondria per cell in WT, MT_long_ and MT_short_ cells normalized to mean total wild-type mitochondrial volume (*n*=10, 12 and 13 cells respectively). In B-D, light grey crosses represent outliers, asterisk represents significance (p<0.05) and ‘n.s.’ indicates no significant difference (one-way ANOVA, Tukey’s Honestly Significant Difference procedure).

We observed that mitochondria underwent increased fission upon microtubule depolymerization, but did not observe their subsequent aggregation as reported previously (Fig. S1A-C). Instead, the fragmented mitochondria were mobile and frequently in close contact with each other (Fig. S1B, C, Movie S1). Since the depolymerization of microtubules had a direct effect on mitochondrial fission, we set out to study the consequence of modification of microtubule dynamics on mitochondrial dynamics. To this end, we visualised the mitochondria and microtubules of fission yeast strains carrying deletions of antagonistic kinesin-like proteins, Klp5/Klp6 and Klp4 in high-resolution deconvolved images (Fig. 1A, Movies S2, S3 and S4).

The heteromeric Klp5/Klp6 motor is required for maintenance of interphase microtubule length by promoting catastrophe at microtubule plus ends (Tischer et al., 2009; West et al., 2001). Cells lacking Klp5 and Klp6 exhibited long microtubule bundles (‘MT_long_’, Fig. 1B) as reported previously due to a decreased catastrophe rate (Tischer et al., 2009). In contrast, Klp4 is required for polarized growth in fission yeast and has been suggested to promote microtubule growth (Browning et al., 2000; Busch et al., 2004). As a result, in the absence of Klp4, microtubule bundles were only about half the length of wild-type bundles (‘MT_short_’, Fig. 1B).

As in wild-type cells, mitochondria in Klp5Δ/Klp6Δ were in close contact with the microtubule, whereas we observed reduced association between the short microtubules and mitochondria in Klp4Δ cells (Fig. 1A, Movies S3 and S4). While wild-type cells had 4.9 ± 0.4 (mean ± s.e.m.) mitochondria per cell, we observed that Klp5Δ/Klp6Δ contained only 2.3 ± 0.4 (mean ± s.e.m.). In contrast, Klp4Δ cells had 10 ± 0.9 mitochondria per cell (mean ± s.e.m., Fig. 1C). This indicated that the number of mitochondria per cell was inversely related to the length of microtubule bundle. However, the decrease in the number of mitochondria in Klp5Δ/Klp6Δ cells and increase in Klp4Δ cells were not at the expense of mitochondrial volume, since the net mitochondrial volume in both cases was comparable to wild-type mitochondrial volume (Fig. 1D), with individual mitochondrial volumes changing to compensate for the difference in mitochondrial numbers between WT, Klp5Δ/Klp6Δ and Klp4Δ cells (Fig. S1D).

### 2. Cells with short microtubules undergo increased fission

To understand the difference in mitochondrial numbers in wild-type, Klp5Δ/Klp6Δ and Klp4Δ cells, we acquired and analyzed time-lapse videos at the single mitochondrion level in all three cases (Fig. 2A, Movies S5, S6 and S7). Similar to our observations from high-resolution images, we measured 3.6 ± 0.2, 2.7 ± 0.2, and 6.9 ± 0.8 mitochondria (mean ± s.e.m.) in wild-type, Klp5Δ/Klp6Δ and Klp4Δ cells respectively (Fig. 1C and 2B). Analysis of evolution of these mitochondrial numbers revealed no significant changes over time (Fig. 2B). Additionally, mitochondria in wild-type cells underwent ~1 fission and ~1 fusion event every minute on an average, whereas Klp5Δ/Klp6Δ cells exhibited a fission frequency that was half that of wild-type, and Klp4Δ mitochondria had a fission frequency that was almost double that of wild-type (Fig. 2C). The fusion frequency of mitochondria in Klp4Δ cells was slightly higher than in wild-type and Klp5Δ/Klp6Δ cells (Fig. 2D), likely due to the increased number of mitochondria in Klp4Δ cells that could participate in fusion. However, the resulting ratio of the mean fission frequency to the mean fusion frequency was ~1, ~0.5 and ~1.3 in wild-type, Klp5Δ/Klp6Δ and Klp4Δ cells respectively. We therefore concluded that the difference in mitochondrial numbers between wild-type cells and Klp5Δ/Klp6Δ and Klp4Δ arose primarily due to the changes in fission frequencies of mitochondria.

**Figure 2.**
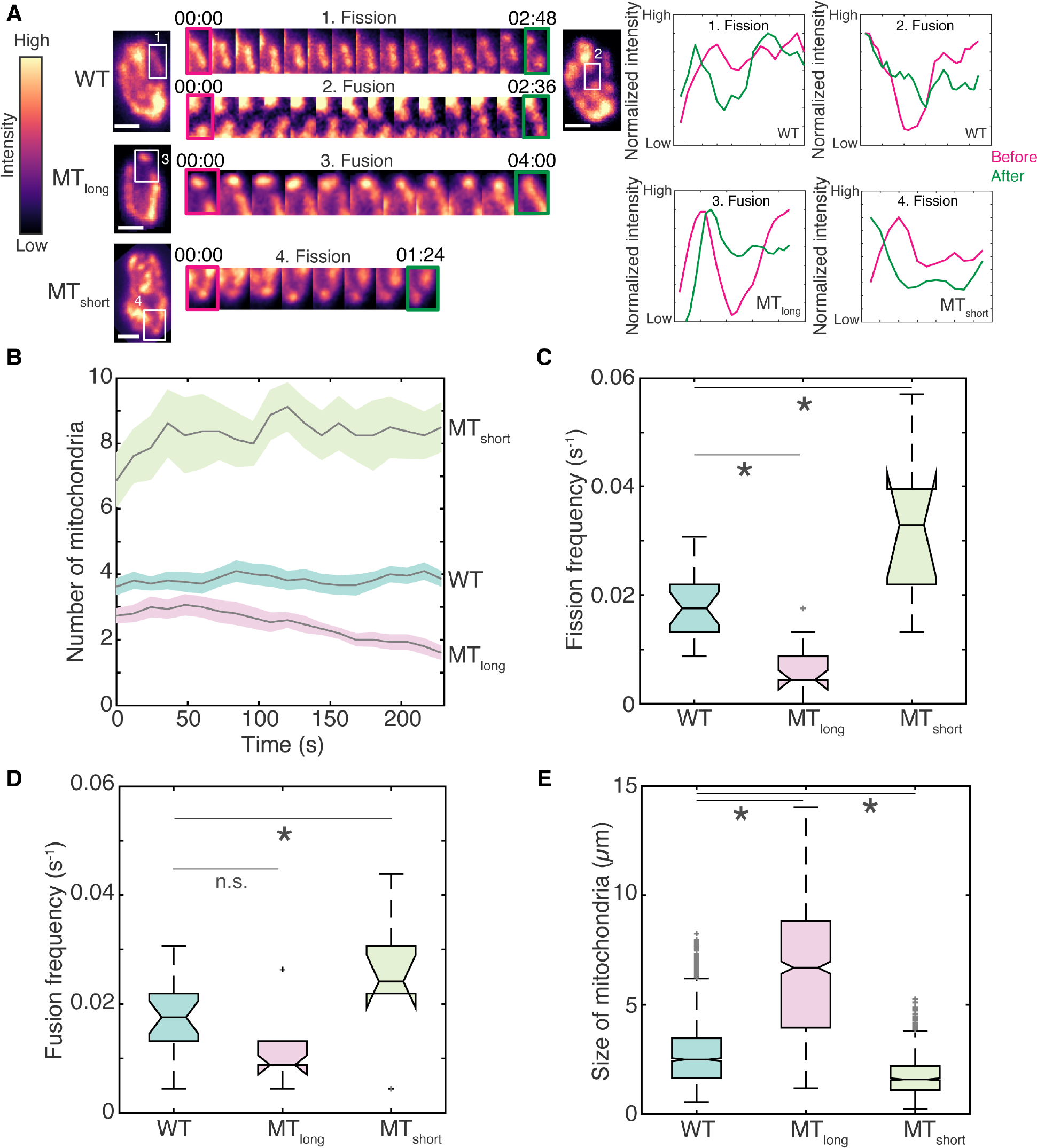
Microtubule length determines fission frequency of mitochondria. **(A)**Montage of maximum intensity projected confocal Z-stack images of wild-type (‘WT’, strain KI001, see Table S1), Klp5Δ/Klp6Δ (‘MT_long_’, strain G3B, see Table S1) and Klp4Δ (‘MT_short_’, strain G5B, see Table S1) cells represented in the intensity map indicated to the left of the images. The insets (white box) and their montages on the right of the images are representative fission and fusion events in WT (‘1’ and ‘2’), fusion event in MT_long_ (‘3’) and fission event in MT_short_ cell (‘4’). Time is indicated as ‘mm:ss’ above the montage of the insets. The normalized intensity along the mitochondrion in the inset before (magenta) and after (green) the fission or fusion event is indicated in plots to the right of the montages. **(B)**Evolution of mitochondrial number over time indicated as mean (solid grey line) and standard error of the mean (shaded region) for WT, MT_long_ and MT_short_ cells (*n*=21, 15 and 8 cells respectively. **(C)**Box plot of the fission frequency of mitochondria per second in WT, MT_long_ and MT_short_ cells (*n*=21, 15 and 8 cells respectively). **(D)**Box plot of the fusion frequency of mitochondria per second in WT, MT_long_ and MT_short_ cells (*n*=21, 15 and 8 cells respectively). **(E)**Box plot of the size of mitochondria in WT, MT_long_ and MT_short_ cells, calculated as the length of the major axis of an ellipse fitted to each mitochondrion (*n*=1613, 739 and 1326 mitochondria respectively). In C-E, light grey crosses represent outliers, asterisk represents significance (p<0.05) and ‘n.s.’ indicates no significant difference (one-way ANOVA in C and D, Kruskal-Wallis test in E, Tukey’s Honestly Significant Difference procedure).

Note that the rate of mitochondrial fission (per second per mitochondrion) was comparable to the rate of mitochondrial fusion (per second per mitochondrion) in all three cases, resulting in the constant number of mitochondria we measured over time (Fig. 2B). As observed in the high resolution deconvolved images, we also measured significant differences in the size and morphology of the mitochondria in WT, Klp5Δ/Klp6Δ and Klp4Δ cells (Fig. 2E, Fig. S2). Mitochondrial sizes reflected microtubule bundle lengths, with the largest mitochondria in Klp5Δ/Klp6Δ cells and the smallest in Klp4Δ cells. WT cells predictably had mitochondrial sizes between that of Klp4Δ and Klp5Δ/Klp6Δ cells (Fig. 2E, Fig. S2).

**Figure 3.**
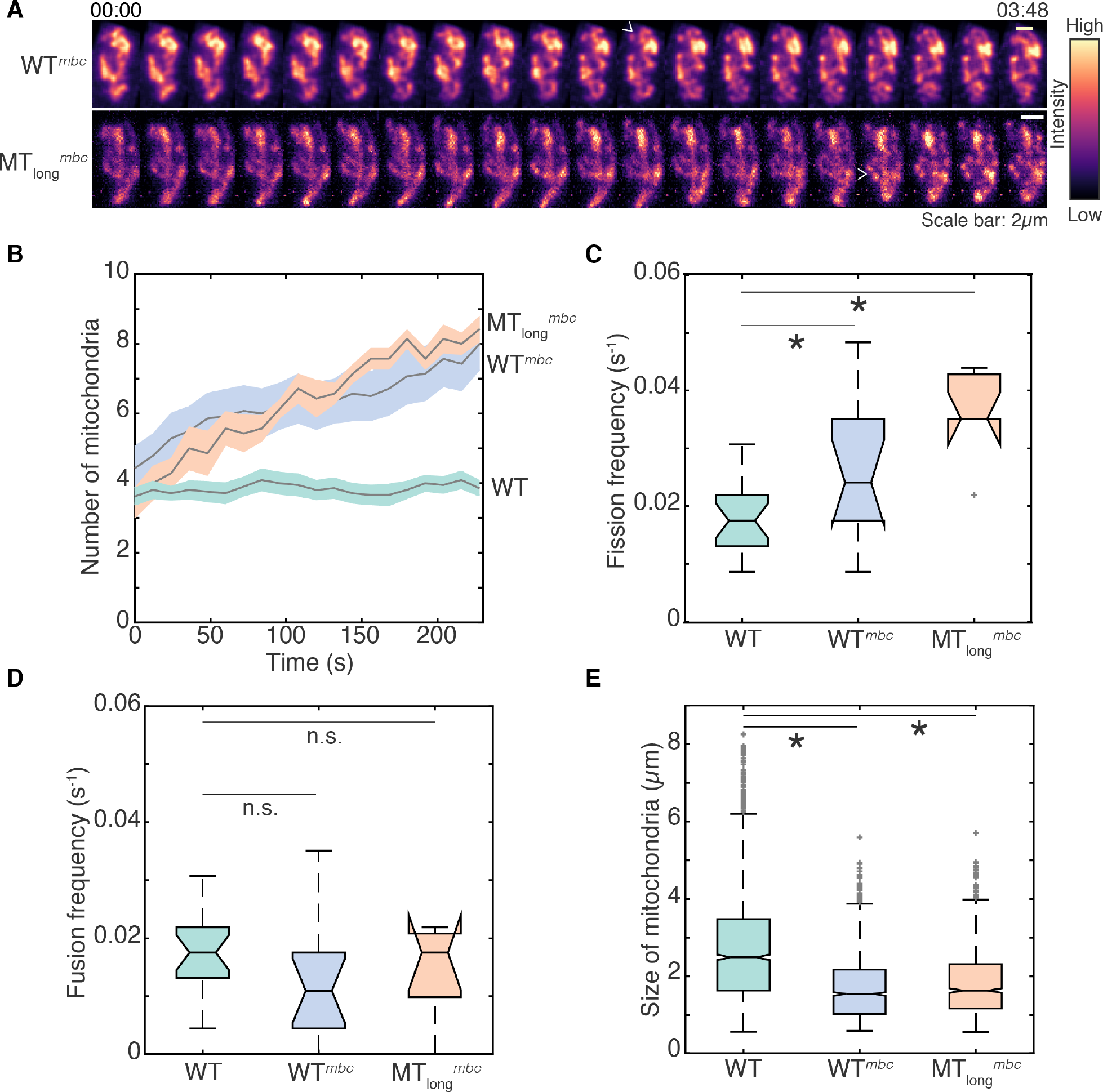
Microtubule depolymerization induces increased fission in WT and Klp5Δ/Klp6Δ cells. **(A)**Montage of maximum intensity projected confocal Z-stack images of MBC-treated wild-type (WT^mbc^, strain KI001, see Table S1) and Klp5Δ/Klp6Δ (‘MT_long_^mbc^’, strain G3B, see Table S1) cells represented in the intensity map indicated to the right of the images. White open arrowheads point to representative fission events. 00:00 indicates time (mm:ss) 2min after addition of MBC. **(B)**Evolution of mitochondrial number with time indicated as mean (solid grey line) and standard error of the mean (shaded region) for wild-type (‘WT’), WT^mbc^, and MT_long_^mbc^ cells (*n*=21, 14 and 7 cells respectively). **(C)**Box plot of the fission frequency of mitochondria per second in WT, WT^mbc^, and MT_long_^mbc^ cells (*n*=21, 14 and 7 cells respectively). **(D)**Box plot of the fusi frequency of mitochondria per second in WT, WT^mbc^, and MT_long_^mbc^ cells (*n*=21, 14 and 7 cells respectively). **(E)**Box plot of the size of mitochondria in WT, WT^mbc^, and MT_long_^mbc^ cells, calculated as the length of the major axis of an ellipse fitted to each mitochondrion (*n*=1613, 1765 and 886 mitochondria respectively). In C-E, light grey crosses represent outliers, asterisk represents significance (p<0.05) and ‘n.s.’ indicates no significant difference (one-way ANOVA in C and D, Kruskal-Wallis Test in E, Tukey’s Honestly Significant Difference procedure). Note that the WT data represented in this figure is the identical to the wild-type data plotted in Fig. 2 and Fig. S2 and has been re-used for comparison.

### 3. Cells devoid of microtubules undergo fission with increased frequency but show unaltered fusion frequencies

While the Klp5Δ/Klp6Δ cells containing long microtubules had the same total volume of mitochondria as WT cells (Fig. 1D), their mitochondria appeared less fragmented. So, to specifically test the role of microtubules in dictating mitochondrial fission, we depolymerized microtubules in wild-type and Klp5Δ/Klp6Δ cells and measured mitochondrial dynamics in time-lapse movies (Fig. 3A, Movies S8, S9). In both WT and Klp5Δ/Klp6Δ cells, upon microtubule depolymerization, we observed a progressive increase in the number of mitochondria (Fig. 3B). We next quantified the fission and fusion events in WT and Klp5Δ/Klp6Δ cells treated with MBC. We measured that the fission frequency of mitochondria in MBC-treated cells was doubled when compared to untreated control cells (Fig. 3C). At the same time, the mitochondrial fusion frequency remained unchanged (Fig. 3D), indicating that the immediate consequence of the loss of microtubules was increased fission, without concomitant changes in mitochondrial fusion. Additionally, upon MBC treatment we measured mitochondrial sizes and morphologies that were reminiscent of Klp4Δ cells (Fig. 3E, Fig. S3A-C).

Increase in oxidative stress via reactive oxygen species (ROS) levels has also been described to induce mitochondrial fission (Pletjushkina et al., 2006). However, we measured no difference in ROS levels between wild-type, Klp5Δ/Klp6Δ, and Klp4Δ cells (Fig. S3D).

### 4. Mitochondria undergo stochastic partitioning during mitosis

We next sought to understand the biological role for increased mitochondrial fission upon microtubule depolymerization. Fission yeast undergoes closed mitosis, wherein the nuclear envelope does not undergo breakdown during cell division (Ding et al., 1997). Upon onset of mitosis in fission yeast, the interphase microtubules that were previously in the cytoplasm are reorganized to form the spindle inside the closed nucleus. This natural situation mimics the depolymerization of microtubules via the chemical inhibitor MBC. Therefore, we set out to study the changes in the mitochondrial network upon cell entry into mitosis. We first obtained high-resolution deconvolved images of the microtubule and mitochondria in fission yeast cells undergoing cell division (Fig. 4A, Fig. S4A, Movie S10). We observed that dividing wild-type cells had ~4× the number of mitochondria in interphase cells (Fig. 4B). Moreover, similar to what was seen in cells lacking microtubules or Klp4 (Fig. 1A), mitochondria in dividing cells were shorter and more rounded (Fig. 4A, Fig. S4A). There was no relationship between length of the mitotic spindle and the number of mitochondria (Fig. 4B, Fig. S4A), indicating that the increased fission likely occurred fairly early upon entry into mitosis. Analysis of time-lapse videos of wild-type cells before and 10min after entry into mitosis revealed a doubling of mitochondrial numbers in this time (Fig. 4C, D, Movie S11). The fragmented mitochondria appeared more mobile and were able to traverse distances of ~1μm in the cell (Fig. 4C). In this same period of time, non-dividing interphase cells did not show any change in mitochondrial numbers (Fig.S4B, C).

**Figure 4.**
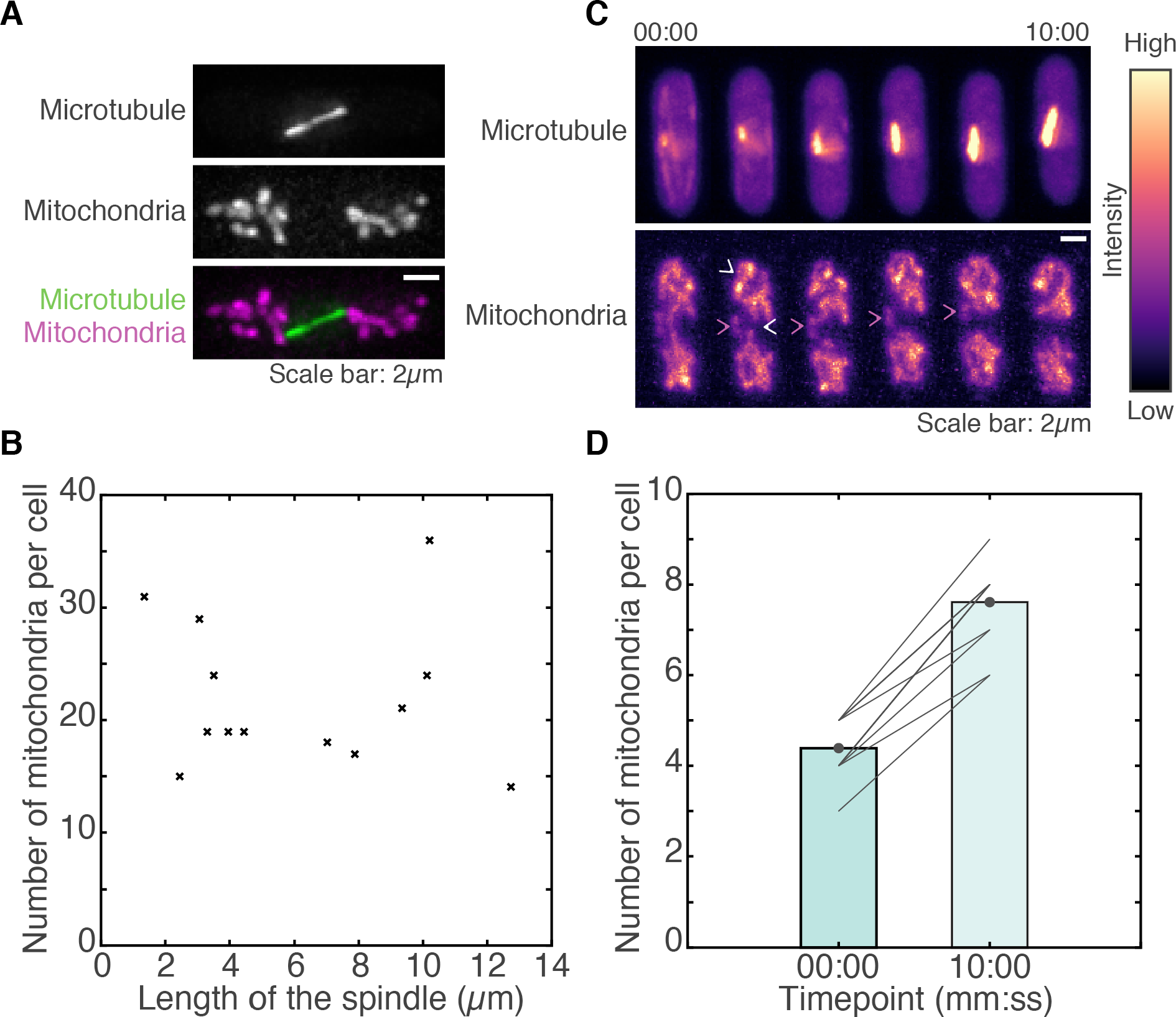
Mitotic cells contain several short mitochondria. **(A)**Maximum intensity projections of deconvolved Z-stack images of the microtubules (top), mitochondria (centre) and their composite (bottom) of a wild-type cell (strain KI001, see Table S1) undergoing division. **(B)**Scatter plot of the length of the mitotic spindle vs. the number of mitochondria per cell in dividing cells (*n*=13 cells). **(C)**Montage of maximum intensity projected confocal Z-stack images of the microtubules (top) and mitochondria (bottom) in a wild-type cell undergoing cell division represented in the intensity map indicated to the right of the images. White open arrowheads point to representative fission events. Magenta arrowheads point to a representative mobile, fragmented mitochondrion. Time is indicated above the images in mm:ss. **(D)**Bar plot of the mean number of mitochondria per cell before (’00:00’) and 10mins after (’10:00’) the onset of mitosis. Solid grey lines represent data from individual cells (*n*=16 cells).

Increase in mitochondrial numbers prior to cell division could aid in increasing the likelihood of equal partitioning of mitochondria between daughter cells, given a stochastic mechanism. To test if mitochondria in our system underwent stochastic, independent segregation (Huh and Paulsson, 2011), with each mitochondrion in the mother cell having a 50% chance of segregating to either of the future daughter cells during mitosis, we tested the fit of our data to a binomial distribution (Birky, 1983) using a Chi-square test as previously described (Hennis and Birky, 1984). Our data (Table S2) did not differ significantly from a binomial distribution with a Chi-square statistic of 7.1846 with 3 degrees of freedom and p=0.0662, indicating that mitochondria in fission yeast were indeed segregating stochastically during cell division. The increase in mitochondrial numbers upon onset of mitosis also served to reduce the partitioning error of mitochondria between the daughter cells as predicted by stochastic segregation (Fig. S4D).

### 5. Cells with long microtubules are protected from unopposed mitochondrial fission

The mitochondrial fission protein in yeast is a dynamin-related GTPase, Dnm1 (Jourdain et al., 2009). Dnm1 brings about the fission of mitochondria by self-assembling into rings or spirals around the mitochondrial outer membrane and employing its GTPase activity to effect the scission (Ingerman et al., 2005). In the absence of Dnm1, mitochondria are organised as extended, fused ‘nets’ (Guillou et al., 2005; Jourdain et al., 2009), which do not undergo fission even in the absence of microtubules (Fig. S5A). Further in Klp4Δ cells, which typically contain several short mitochondria (Fig. 1A), absence of Dnm1 results in a single large, fused mitochondrion (Fig. S5B). Therefore, all mitochondrial fission in *S. pombe* is reliant on the activity of Dnm1. Additionally, during mitosis, cells lacking Dnm1 that contained a single large mitochondrion relied on the cytokinesis of the mother cell to also split the mitochondrion into the daughter cells (Fig. S5C).

**Figure 5.**
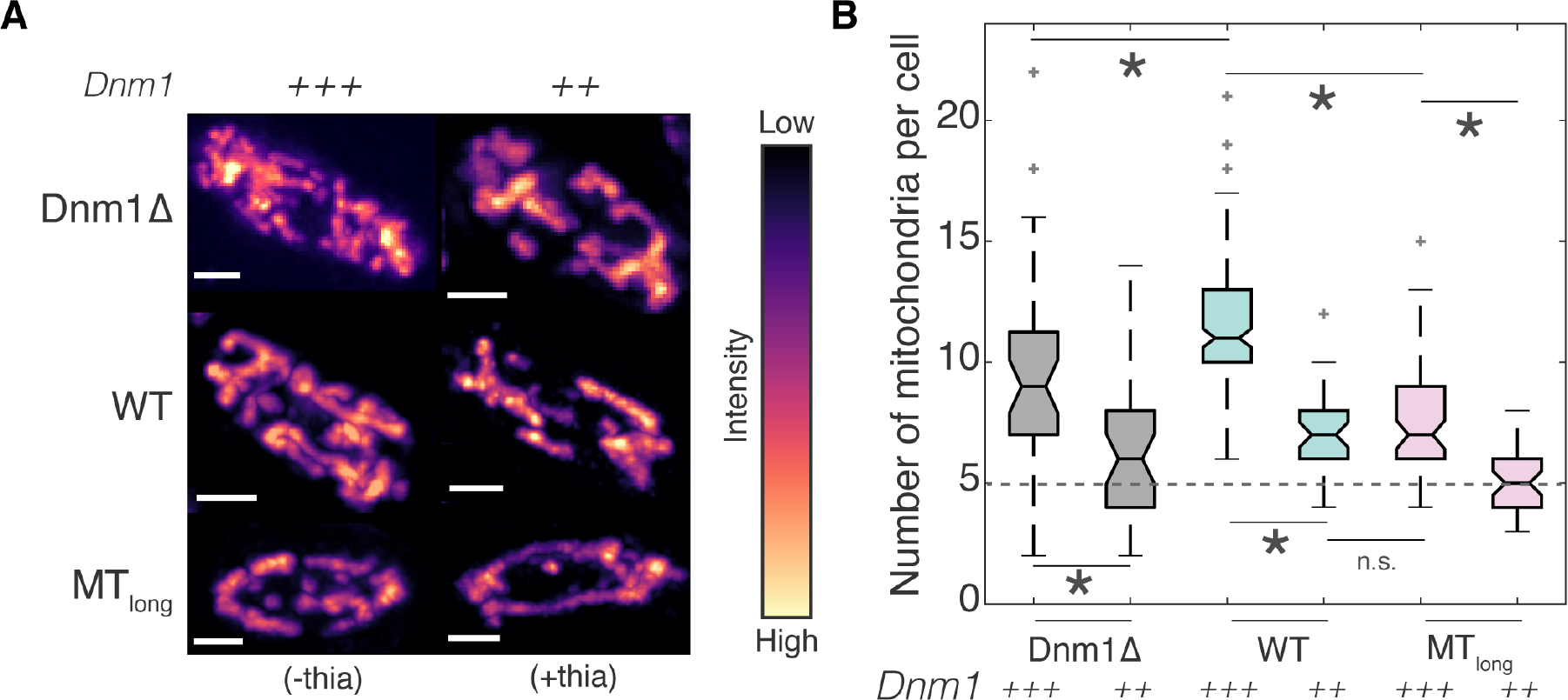
Association with microtubules prevents unopposed fission of mitochondria. **(A)**Maximum intensity projections of deconvolved Z-stack images of mitochondria in cells transformed with functional untagged Dnm1 (plasmid pREP41-Dnm1, see Table S1) in Dnm1Δ cells (see Table S1), wild-type (‘WT’, strain FY7143, see Table S1) and Klp5Δ/Klp6Δ cells (‘MT_long_’, strain FY20823, see Table S1) represented in the intensity map indicated to the right of the images. High overexpression of Dnm1 is indicated with ‘*+++*’ (‘(-thia)’) and low overexpression with ‘*++*’ (‘+thia)’). Scale bars represent 2μm**. (B)**Box plot of mitochondrial numbers in Dnm1Δ, WT and MT_long_ cells with high overexpression (‘*+++*’) or low expression of Dnm1 (‘*++*’) (*n*= 37, 36, 149, 37, 63, 41 cells respectively). The mean mitochondrial number in WT cells expressing normal amount of Dnm1 is depicted by the dashed line. Light grey crosses represent outliers, asterisk represents significance (p<0.05) and ‘n.s.’ indicates no significant difference (Kruskal-Wallis test, Tukey’s Honestly Significant Difference Procedure).

Taken together, our results suggest that association of mitochondria with microtubules inhibits mitochondrial fission. Previously, cryo-electron tomographic analysis in fission yeast indicated a preferred separation distance of ~20nm for mitochondria associated with microtubules (Höög et al., 2007). So too, cryo-elctron microscopy in budding yeast revealed that Dnm1 assembled into rings with an outer diameter of 129nm and lumen diameter of 89nm, resulting in ~20nm-high structures around mitochondria (Mears et al., 2011). Therefore, mitochondria associated with microtubules might have insufficient space to accommodate the Dnm1 ring around their diameter, thereby inhibiting fission due to simple physical constraints. To test this hypothesis, we first transformed Dnm1Δ, wild-type, and Klp5Δ/Klp6Δ cells with a plasmid expressing Dnm1 under the control of the nmt1 promoter (see Table S1) and visualized the mitochondria (Fig. 5A). These cells exhibit high overexpression of Dnm1 in the absence of thiamine, and low overexpression in the presence of 0.05μM thiamine in the culture medium. We counted the number of mitochondria present in these cells and estimated that Dnm1Δ cells contained 9.5 ± 0.7 and 6.4 ± 0.5 mitochondria (mean ± s.e.m) with high overexpression and low overexpression of Dnm1 respectively (Fig. 5B). Wild-type cells highly overexpressing Dnm1 had 11.6 ± mitochondria (mean ± s.e.m), which is twice that of wild-type cells expressing normal levels of Dnm1 (Fig. 5B). The increase in mitochondrial numbers in these cells is due to the increase in Dnm1 numbers, that likely accelerates the kinetics of the Dnm1 ring formation and hence, mitochondrial fission. In wild-type cells that were grown in the presence of thiamine, the mitochondrial number was 7 ± 0.3 (mean ± s.e.m.), presumably because of low overexpression of Dnm1.

**Figure 6.**
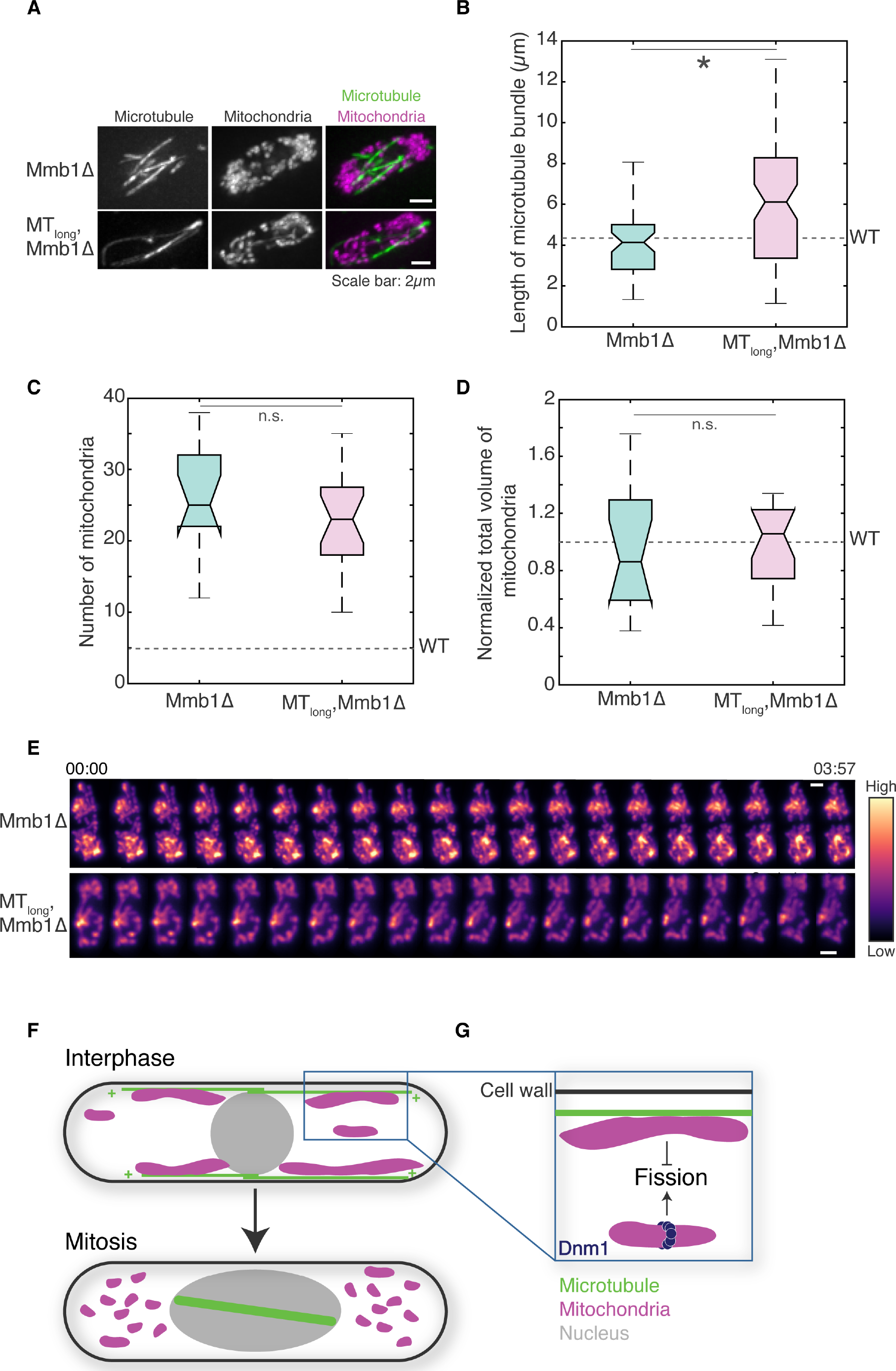
Deletion of Mmb1 results in unopposed fission in WT and MT_long_ cells. **(A)**Maximum intensity projections of deconvolved Z-stack images of microtubules (left), mitochondria (centre) and their composite (right) in Mmb1Δ cells (top, strain PT2244, see Table S1) and Klp5Δ/Klp6Δ-Mmb1Δ cells (‘MT_long_, Mmb1Δ’, bottom, strain VA076 see Table S1). **(B)**Box plot of microtubule bundle length in Mmb1Δ (*n*= 54 microtubules from 13 cells) and Klp5Δ/Klp6Δ-Mmb1Δ cells (*n*=81 microtubules from 25 cells respectively). The mean microtubule bundle length in WT cells is depicted by the dashed line. **(C)**Box plot of mitochondrial numbers in Mmb1Δ and Klp5Δ/Klp6Δ-Mmb1Δ cells (*n*=14 and 20 cells respectively). The mean mitochondrial number in WT cells is depicted by the dashed line. **(D)**Box plot of mitochondrial volume in Mmb1Δ and Klp5Δ/Klp6Δ-Mmb1Δ cells (*n*=14 and 20 cells respectively) normalized to mean total mitochondrial volume in WT cells. The mean mitochondrial volume in WT cells is depicted by the dashed line. **(E)**Montage of maximum intensity projected confocal Z-stack images of the mitochondria in a Mmb1Δ cell (top) and Klp5Δ/Klp6Δ-Mmb1Δ cell (‘MT_long_, Mmb1Δ’, bottom). The intensity map is indicated to the right of the images. Time is indicated above the images in mm:ss. In B-D, asterisk represents significance (p<0.05), ‘n.s.’ represents no significant difference (one-way ANOVA, Tukey’s Honestly Significant Difference Procedure). **(F)**Model of mitochondrial dynamics mediated by microtubule dynamics. Microtubules polymerize and depolymerize at their plus ends (‘+’). Absence of microtubule bundles in the cytoplasm during cell division enables the fragmentation of mitochondria. **(G)**When mitochondria are bound to microtubules, Dnm1 assembly is inhibited. Upon microtubule depolymerization, this inhibition is alleviated and Dnm1 can effectively mediate scission of mitochondria.

Interestingly, in Klp5Δ/Klp6Δ cells with high and low overexpression of Dnm1, we counted only 7.5 ± 0.3 mitochondria and 5.2 ± 0.2 mitochondria (mean ± s.e.m.) respectively. In Klp5Δ/Klp6Δ, the presence of longer microtubules than wild-type cells possibly prevented increased fission of mitochondria even when cells overexpressed Dnm1. In fact, in Klp5Δ/Klp6Δ cells where there was low overexpression of Dnm1, the mitochondrial number was comparable to that in wild-type cells expressing normal levels of Dnm1 (Fig. 5B).

We additionally employed cells expressing mCherry-tagged microtubules, GFP-tagged Dnm1 and Mitotracker-stained mitochondria and visualized the localization of Dnm1 with respect to the microtubules and mitochondria in the same cells (Fig. S6A). While 97.5% of Dnm1 spots colocalized with the mitochondria that were not bound to the microtubule (115 out of 118 spots, n=18 cells), only 10.2% (12 out of 118 spots, n=18 cells) localized on mitochondria that were attached to the microtubule. While these results indicated an inhibitory role for microtubule-mitochondrial interactions in Dnm1 assembly, the non-functionality of fluorescently-tagged Dnm1, that has been documented in other literature (Jourdain et al., 2009), prevented direct visualization of fission of mitochondria by Dnm1 in these cells.

### 6. Dissociation of mitochondria from microtubules leads to unopposed fission

To further test the hypothesis that Dnm1 assembly was physically impeded on microtubule-bound mitochondria, we sought to move microtubules and mitochondria apart without perturbing the microtubule cytoskeleton. To this end, we employed cells devoid of the protein that links mitochondria with microtubules in fission yeast, Mmb1 (Fig. 6A, Movies S12, S13). First, we measured microtubule bundle lengths in Mmb1Δ cells and Klp5Δ/Klp6Δ-Mmb1Δ cells. We did not observe a significant difference in microtubule bundle lengths between WT and Mmb1Δ cells (Fig. 6B), but Klp5Δ/Klp6Δ-Mmb1Δ cells exhibited microtubules that were significantly longer than WT and Mmb1Δ cells, comparable to Klp5Δ/Klp6Δ cells.

Next, we counted 26.2 ± 1.8 (mean ± s.e.m.) mitochondria in these cells, which is significantly higher than that in WT cells (Fig. 6C). In the case of Klp5Δ/Klp6Δ-Mmb1Δ cells, although microtubules are significantly longer than in WT cells (Fig. 6B), the absence of the linker between microtubules and mitochondria resulted in extensive fission of mitochondria.

We counted 23.3 ± 1.4 (mean ± s.e.m.) mitochondria in Klp5Δ/Klp6Δ cells lacking Mmb1Δ (Fig. 6C), which was not significantly different from Mmb1Δ cells. Additionally, the total mitochondrial volume in both Mmb1Δ and Mmb1Δ-Klp5Δ/Klp6Δ cells was also unchanged compared to WT cells indicating a change in morphology of mitochondria, but not biogenesis (Fig. 6D). Time-lapse images of mitochondria in Mmb1Δ and Mmb1Δ-Klp5Δ/Klp6Δ cells revealed mobile mitochondrial fragments, consistent with the movement expected for mitochondria that are not bound to microtubules (Fig. 6E). Therefore, by separating mitochondria from microtubules, we alleviated the inhibition of Dnm1 assembly on the mitochondria, thereby promoting mitochondrial fission.

Fig. S6B contains a summary of the mitochondrial parameters measured in this study.

## Discussion

We discovered that association with microtubules determines mitochondrial dynamics and thereby, mitochondrial morphology (Fig. 6F). While previous studies discounted the role of motors in the determination of mitochondrial positioning in fission yeast (Brazer et al., 2000; Chiron et al., 2008; Li et al., 2015; Yaffe et al., 2003), we have identified kinesin-like proteins that regulate mitochondrial morphology through their control over microtubule length. The association of mitochondria with microtubules mediated by the protein Mmb1 was found to be necessary and sufficient to maintain equilibrium between fission and fusion of mitochondria, with loss of Mmb1 leading to unopposed fission.

In mammalian cells, mitochondria employ microtubule-associated machinery primarily to traverse the cell. However, mitochondrial dynamics have been observed to be related to microtubule-assisted processes. For instance, in HeLa cells, Drp1 recruitment to the mitochondria was observed to be dependent on the motor protein cytoplasmic dynein (Varadi et al., 2004). Mobile mitochondria on microtubules have also been seen to participate in kiss-and-run fusions that are transient (Liu et al., 2009). However, a direct role for the microtubule cytoskeleton in the maintenance of mitochondrial dynamics has not yet been explored.

We observed that fission yeast cells fragment mitochondria by emptying the cytoplasm of its microtubules upon onset of mitosis, thereby ensuring stochastic partitioning (Birky, 1983) of the several small mitochondria between future daughter cell. The increase in mitochondrial numbers prior to cell division likely serves to create a well-mixed, homogeneous cytoplasm that is primed for stochastic segregation of mitochondria (Fig. 6F). Mitochondrial fragmentation also reduces the partitioning error between the daughter cells (Hennis and Birky, 1984), since mitochondrial numbers of 5 or less, which is typical of wild-type interphase cells, exhibit errors between ~45 and 100% (Fig. S4D). While mammalian cells also exhibit fragmented mitochondria during mitosis (Mitra et al., 2009; Taguchi et al., 2007), it has not yet been conclusively demonstrated that segregation to daughter cells is achieved by stochastic mechanisms. As we observed here, in cells containing mutants of the mitochondrial fission protein, mammalian cells employ the cytokinesis machinery to partition mitochondria between daughter cells (Ishihara et al., 2009). However, this mechanism of mitochondrial partitioning in Drp1-mutant cells is less uniform since mitochondria exist as a large network at the time of division (Mishra and Chan, 2014).

We observed that the presence of long microtubules was inhibitory for fission of mitochondria even while Dnm1 was overexpressed. Wild-type cells contained the same total volume of mitochondria as cells with long microtubules, but exhibited increased mitochondrial fission in the presence of increased Dnm1 levels. In disease states such as cancer (Ferreira-da-Silva et al., 2015) and neurodegeneration (Itoh et al., 2013), mitochondrial fragmentation has been linked with overexpression of Drp1 in mammalian cells. In this work, we have identified that cells that present increased microtubule stabilization are able to overcome the effect of Dnm1 overexpression.

By dissociating mitochondria from microtubules without perturbing the microtubule cytoskeleton, we demonstrated that the failure of Dnm1 ring assembly in the presence of microtubules was the nexus between microtubule dynamics and mitochondrial dynamics (Fig. 6G). This finding could have interesting implications for the interactions between microtubules and mitochondria in other cell types, including in neurons where at any given time, a majority of the mitochondria are seemingly stably attached to the microtubule cytoskeleton (Kang et al., 2008). Future studies in these cells will illuminate the consequence of microtubule-binding on mitochondrial dynamics.

Alternatively, increased fission upon microtubule depolymerization might be brought about by phosphorylation of the mitochondrial fission protein Dnm1. Mammalian Drp1 undergoes phosphorylation at S616 mediated by Cdk1/cyclin B during cell division and increases mitochondrial fission (Taguchi et al., 2007). However, fission yeast Dnm1 however lacks this phosphorylation site. Drp1 also contains another phosphorylation site at S637, which is conserved in Dnm1. However, PKA-mediated phosphorylation at S637 has been shown to inhibit fission by Drp1 (Chang and Blackstone, 2007), whereas Ca++/calmodulin-dependent protein kinase α-dependent phosphorylation of Drp1 S637 in rat hippocampal neurons was shown to increase fission (Han et al., 2008). To date, yeast Dnm1 has not been demonstrated to undergo phosphorylation at this site.

Another possibility is that there exists that an unidentified Dnm1-activating factor that is sequestered by microtubules. This could explain the increase in mitochondrial fragmentation upon microtubule depolymerization. However, in our experiments with cells lacking Mmb1, we induced mitochondrial fragmentation by dissociating mitochondria from microtubules without affecting the microtubule cytoskeleton. Therefore, it is unlikely that mitochondrial fission is promoted by a factor released upon microtubule depolymerization. In conclusion, we have discovered a novel mechanism of regulation of mitochondrial fission by the physical association of mitochondria with microtubules.

## Acknowledgments

We thank J. M. Thankachan, S. S. Nuthalapati and M. Ayushman for pilot experiments; High Content Imaging Facility at BSSE, IISc, and P. I. Rajyaguru for the use of the InCell 6000, and Deltavision RT microscopes respectively; P. Delivani, I. Jourdain, R.C. Salas, M. Takaine, I. Tolic, Y. Gachet, P. Tran, and NBRP Japan for yeast strains and constructs; S. Jain for CellROX reagent. The research was supported by the Department of Science and Technology (India)-INSPIRE Faculty Award, the Department of Biotechnology (India) Innovative Young Biotechnologist Award, and the Science and Engineering Research Board (SERB, India) Early Career Research Award awarded to V.A., and SERB Early Career Research Award awarded to S.J.

## Author contributions

K.M. and L.A.C carried out and analyzed the Dnm1 and Mmb1 experiments, M.K.C. and S. J. carried out and analyzed the flow cytometry experiments, V.A. performed all the other experiments and data analysis, designed the project, and wrote the paper.

## Supplementary Materials and Methods

### ROS detection and viability assay using flow cytometry

A loopful of L972, FY20823 or McI438 cells were inoculated in liquid YE and incubated overnight at 30°C in a shaking incubator. Cells were harvested using a table-top centrifuge (2000g, 30s) and washed three times with 1X PBS. The pellets were re-suspended in 500μl of 1X PBS, containing CellROX (5μM) dye (ThermoFisher, USA). Following incubation at 30°C for 30min, cells were washed three times with 1X PBS. Finally, cell pellets were re-suspended in an appropriate volume of 1X PBS containing propidium iodide (6μg/ml), incubated for 10min at room temperature, and run on a flow cytometer (BD FACS-Celesta). A minimum of 10,000 events of cells was collected, and data analyzed using FlowJo (FlowJo LLC, USA). ROS was reported as median fluorescence intensity for each dataset.

### Estimation of partitioning error during cell division

The partitioning error of mitochondria at cell division between the daughter cells *Q*_*x*_ was calculated from the equation (Huh and Paulsson, 2011):

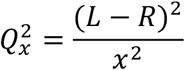

where *L* and *R* were mitochondrial numbers in the resulting daughter cells and *x*, the number of mitochondria in the mother cell. The data in Fig. S4D were simulated by independently assigning *x* mitochondria with 50% probability to either daughter cell. Simulations were re-run 50 times to obtain the mean and the standard error of the mean depicted in Fig. S4D.

### Counting of Dnm1 spots

Dnm1 was visualised in cells grown in the presence of 0.05μM thiamine for partial induction of the nmt1 promoter. Even with only partial induction, a majority of the cells showed extremely high expression of Dnm1, with most of the signal concentrated in a single spot in the cell. The rest of the cells showed lower expression, with Dnm1 signal distributed in a few spots across the cell. The latter were chosen for analysis of number of Dnm1 spots. For counting Dnm1 spots, the maximum intensity projection of deconvolved Z-stack images were used. For identification of spots, we set the lower threshold as 60-80% of the maximum intensity in these images. The contrast in the images in Fig. S6B reflect these threshold settings.

### Supplementary Tables

**Table S1.** Strains and constructs used in this study

| Name | Phenotype | Source |
| --- | --- | --- |
| Dnm1Δ | *h- dnm1::kanr leu1-32ade-* | Yannick Gachet, Toulouse |
| FY20823 | *h- leu1 ura4 his7 Δklp5::ura4+ Δklp6::ura4+* | NBRP, Japan |
| FY7143 | *h- ura4-D18 leu1-32 ade6-M216 his7-366* | Iva Tolić, Croatia |
| G3B | *h- Δklp5- Δklp6 – nmt1-GFP-atb2 leu ade* | Rafael Carazo Salas, UK |
| G5B | *h- klp4::kanr nmt1-GFP-atb2 leu ura* | Rafael Carazo Salas, UK |
| KI001 | *h+ sid4-GFP::kanr kanr -nmtP3-GFP-atb2+ nmt1-pCOX4RFP::leu1+ ura4-D18 ade6-M210* | Iva Tolić, Croatia |
| L972 | *h- WT* | Iva Tolić, Croatia |
| Mcl438 | *h+ tea2d:his3 ade6 leu1-32 ura4-D18 his3-D1* | Iva Tolić, Croatia |
| MTY271 | *h- mCherry-atb2:hphMX6 leu1-32 ura-d18* | Masakatsu Takaine, Japan |
| PT2244 | *h+ mmb1Δ:Kanr cox4-GFP:leu2 mCherry-atb2:Hygr ade6-m210 leu1-32 ura4-d18* | Phong Tran, USA |
| PT1650 | *h+ cox4-GFP:leu1 ade6-M210 leu1-32 ura4-D18* | Phong Tran, USA |
| VA076 | *h+ mmb1Δ:Kanr cox4-GFP:leu2 mCherry-atb2:Hygr Δklp5::ura4+ Δklp6::ura4+ ade6-m210 his7* | This study |
| VA084 | *h- dnm1::kanr klp4::kanr nmt1-GFP-atb2 leu1-32 ade-* | This study |
| FYP2026 (plasmid) | *nmt-GFP-atb2: LEU2* | NBRP, Japan |
| pREP41-Dnm1 (plasmid) | *Dnm1 (untagged)* | Isabelle Jourdain, UK |
| pREP41-Dnm1-Cterm-GFP (plasmid) | *Dnm1-GFP* | Isabelle Jourdain, UK |

**Table S2.**
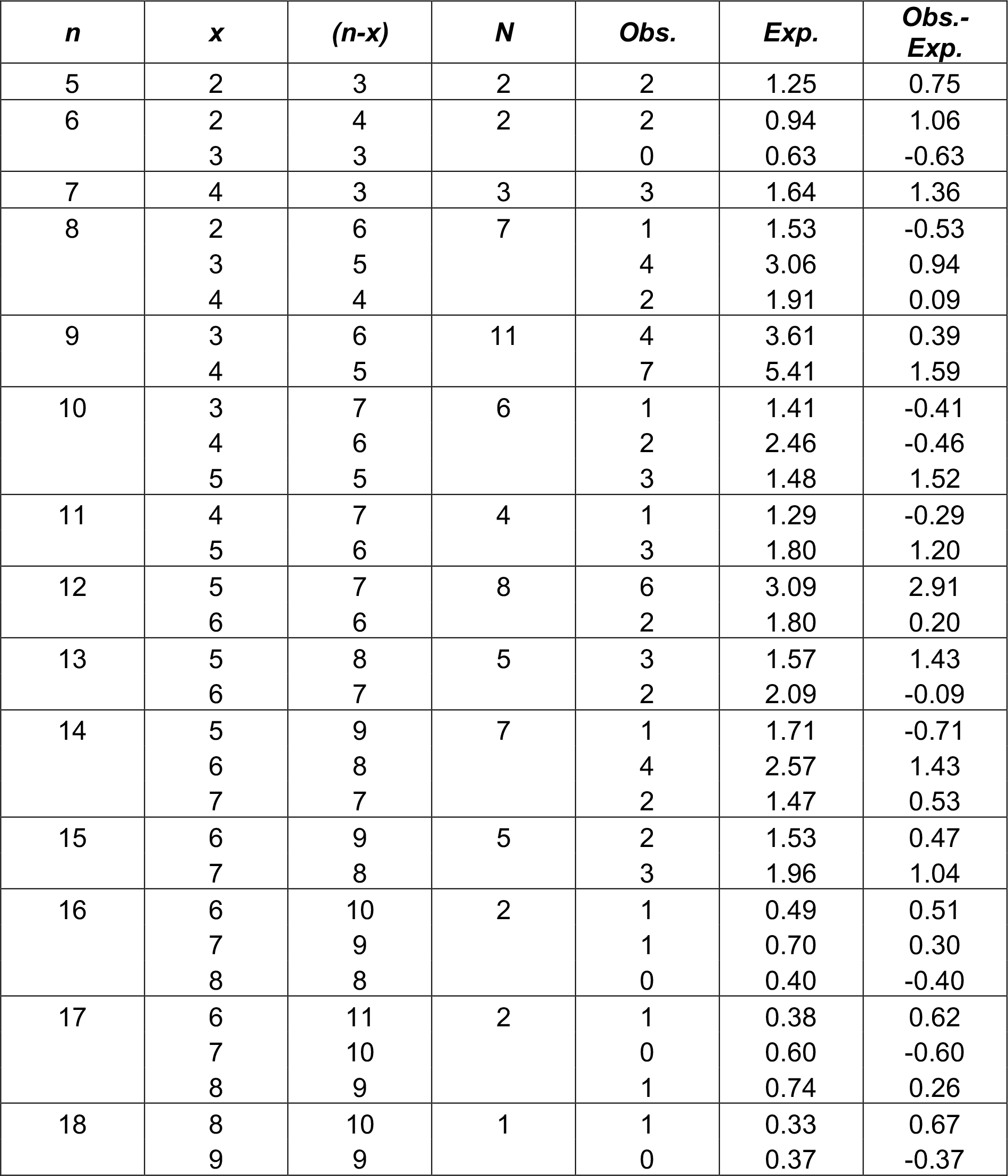
Partitioning of mitochondria between daughter cells. *n* is the number of mitochondria in the mother, *x* and *n-x* in the daughter cells at cell division. *N* is the number of cells counted with *n* mitochondria in the mother. *Obs.* is the observed number of cells showing partition of *n* into *x* and *n-x*, and *Exp.* is the expected number of cells with the same *x* and *n-x* partition given *N* cells predicted by the binomial distribution.

### Supplementary Figures

**Figure S1.**
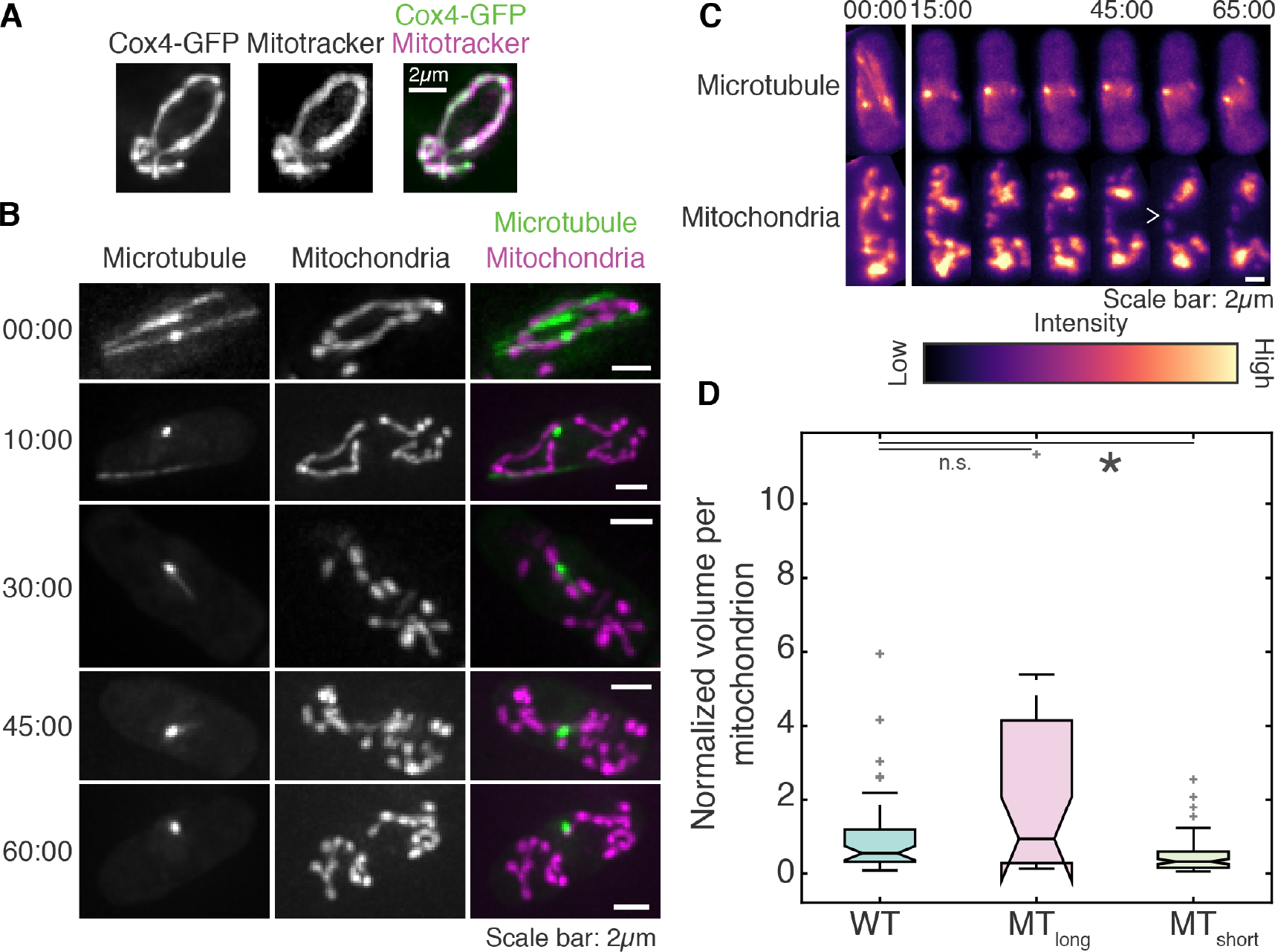
Depolymerization of microtubules increases mitochondrial fission. **(A)**Maximum intensity projections of deconvolved Z-stack images of Cox4-GFP (left), Mitotracker Orange staining (centre) and their composite (right) in interphase cells (strain PT1650, see Table S1) showing colocalization of both signal intensities. **(B)**Maximum intensity projections of deconvolved Z-stack images of microtubules (left), mitochondria (centre) and their composite (right) in fission yeast cells treated with MBC (strain KI001, see Methods). Time in mm:ss is indicated to the left of the images. **(C)**Montage of maximum intensity projected confocal Z-stack images of microtubule depolymerization (top) and mitochondria (bottom) of wild-type cells (strain KI001, see Table S1) treated with MBC (see Methods) represented in the intensity map indicated at the bottom of the images. The open white arrowhead indicates a representative fission event. **(D)**Box plot of the volume of individual mitochondria in wild-type (‘WT’), Klp5Δ/Klp6Δ (‘MT_long_’) and Klp4Δ (‘MT_short_’) cells normalized to mean mitochondrial volume in wild-type cells (*n*=49, 28 and 100 mitochondria respectively). Light grey crosses represent outliers, asterisk represents significance (p<0.05) and ‘n.s.’ indicates no significant difference (Kruskal-Wallis test, Tukey’s Honestly Significant Difference Procedure).

**Figure S2.**
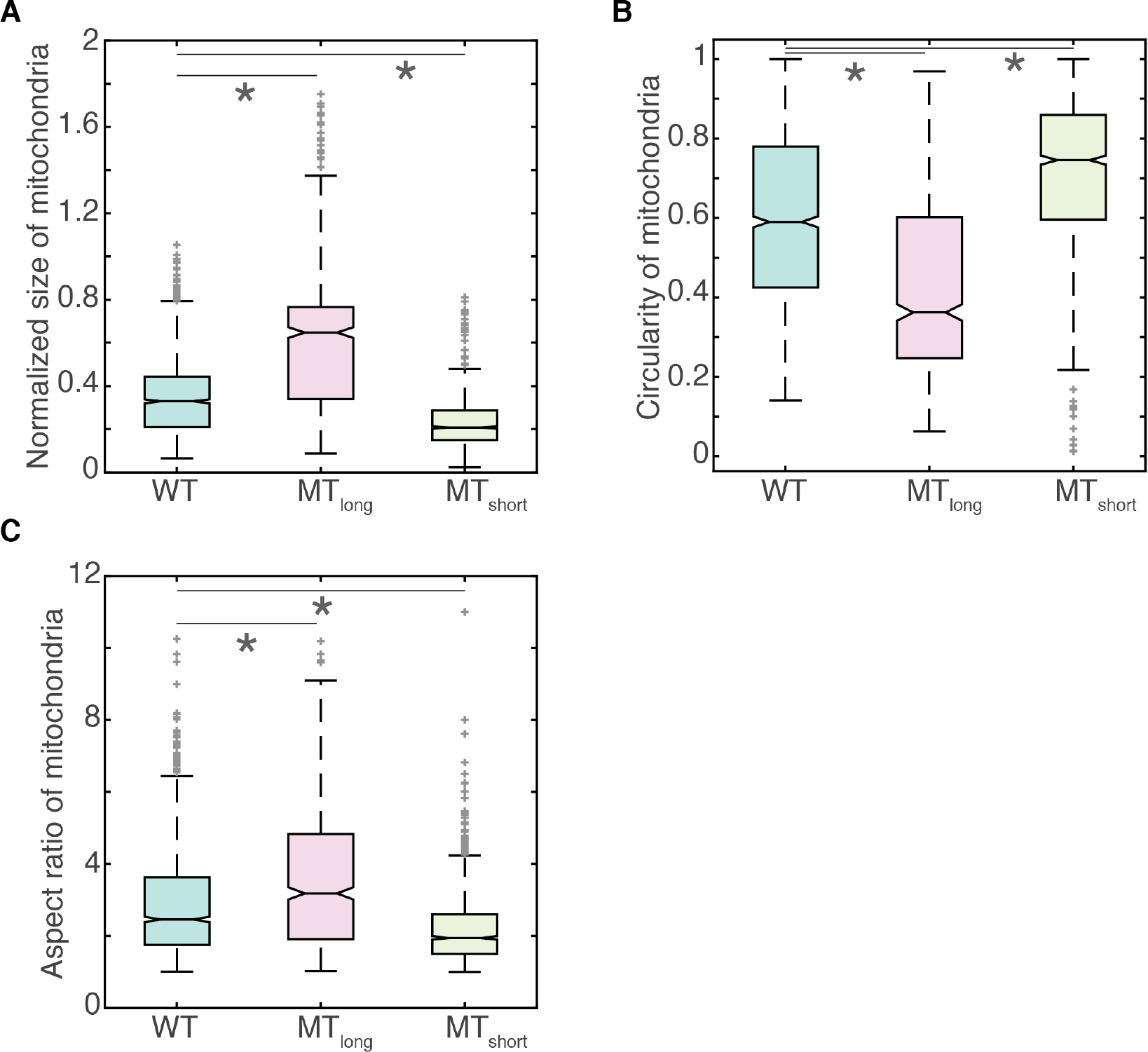
Microtubule lengths determine the morphology of mitochondria. **(A)**Box plot of the size of individual mitochondria in wild-type (‘WT’), Klp5Δ/Klp6Δ (‘MT_long_’) and Klp4Δ (‘MT_short_’) cells normalized to their cell lengths (*n*=1613, 739 and 1326 mitochondria respectively). **(B)**Box plot of the circularity of mitochondria in WT, MT_long_ and MT_short_ cells (*n*=1613, 739 and 1326 mitochondria respectively). **(C)**Box plot of the aspect ratio of mitochondria in WT, MT_long_ and MT_short_ cells (*n*=1613, 739 and 1326 mitochondria respectively). In A-C, light grey crosses represent outliers, asterisk represents significance (p<0.05) and ‘n.s.’ indicates no significant difference (Kruskal-Wallis test, Tukey’s Honestly Significant Difference Procedure).

**Figure S3.**
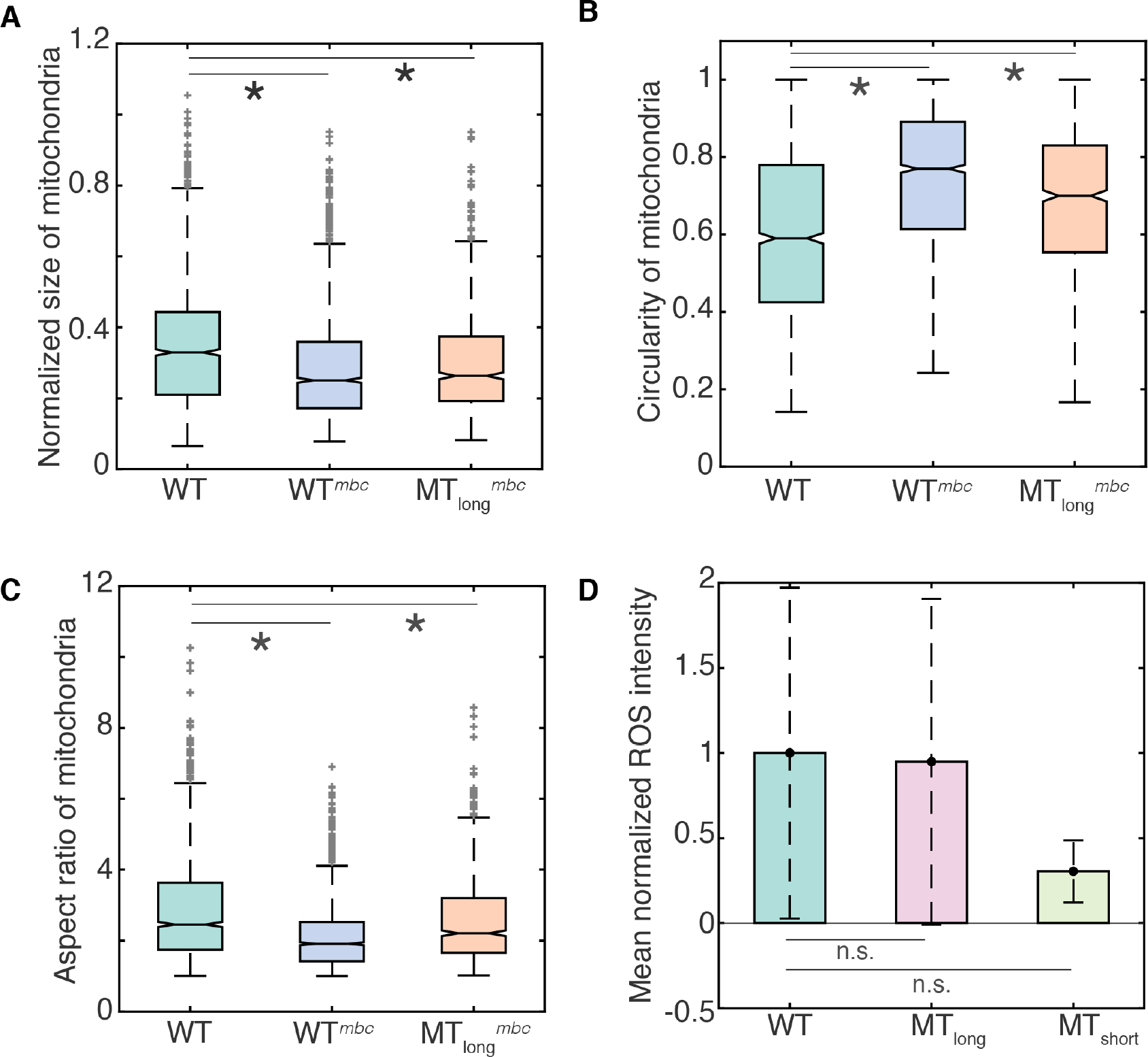
Morphology of mitochondria is altered by treatment with MBC. **(A)**Box plot of the size of individual mitochondria in WT, WT^mbc^, and MT_long_^mbc^ cells normalized to their cell lengths (*n*=1613, 1765 and 886 mitochondria respectively). **(B)**Box plot of the circularity of mitochondria in WT, WT^mbc^, and MT_long_^mbc^ cells (*n*=1613, 1765 and 886 mitochondria respectively). **(C)**Box plot of the aspect ratio of mitochondria in WT, WT^mbc^, and MT_long_^mbc^ cells (*n*=1613, 1765 and 886 mitochondria respectively). **(D)**Bar plot of cellular ROS measured using flow cytometry (see Supplementary Information) in live WT, MT_long_ and MT_short_ cells normalized to mean of the ROS intensity in WT cells (*n*=3 independent experiments, ≥10,000 events each). Error bars represent standard deviation. In A-D, light grey crosses represent outliers, asterisk represents significance (p<0.05) and ‘n.s.’ indicates no significant difference (Kruskal-Wallis test in A-C, one-way ANOVA in D, Tukey’s Honestly Significant Difference Procedure). Note that the WT data represented in this figure is the identical to the wild-type data plotted in Fig. S2 and has been re-used for comparison.

**Figure S4.**
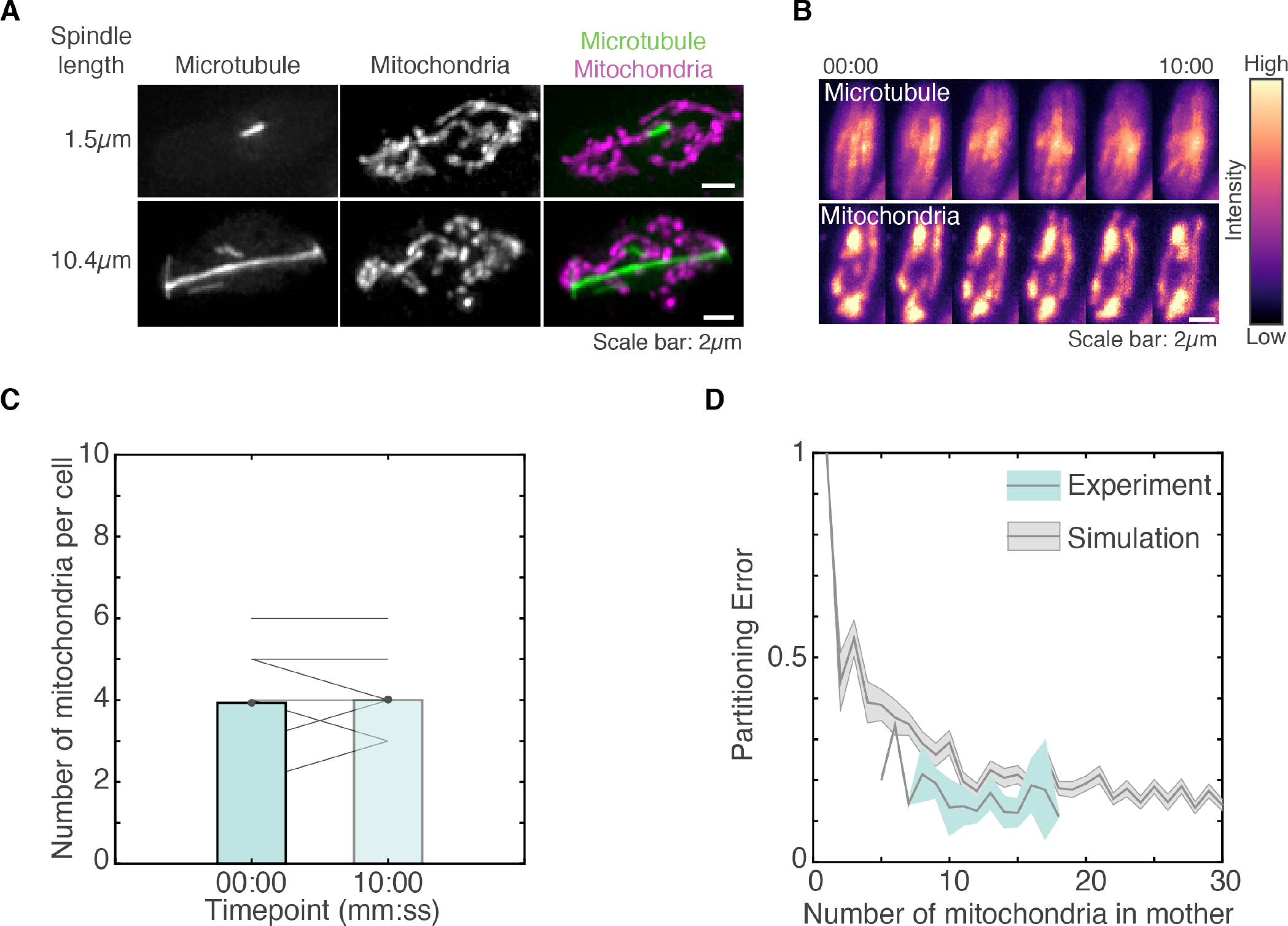
Mitotic spindle length does not influence number of mitochondria. **(A)**Maximum intensity projections of deconvolved Z-stack images of the microtubules (left), mitochondria (centre) and their composite (right) of WT cells (strain KI001, see Table S1) with short (top) and long (bottom) mitotic spindles. The length of the spindle is indicated to the left of the images. **(B)**Montage of maximum intensity projected confocal Z-stack images of the microtubules (top) and mitochondria (bottom) in a non-mitotic wild-type cell (strain KI001, see Table S1) represented in the intensity map indicated to the right of the images. **(C)**Bar plot of the mean number of mitochondria per cell before (’00:00’) and after (’10:00’) the same time window for which mitotic cells were monitored in Fig. 3C, d. Solid grey lines represent data from individual cells (*n*=7 cells). **(D)**Plot of number of mitochondria in mother cell vs. partitioning error (see Supplementary Information) between daughter cells in simulation (grey) and experiments (blue) expressed as mean (solid grey line) and standard error of the mean (shaded area).

**Figure S5.**
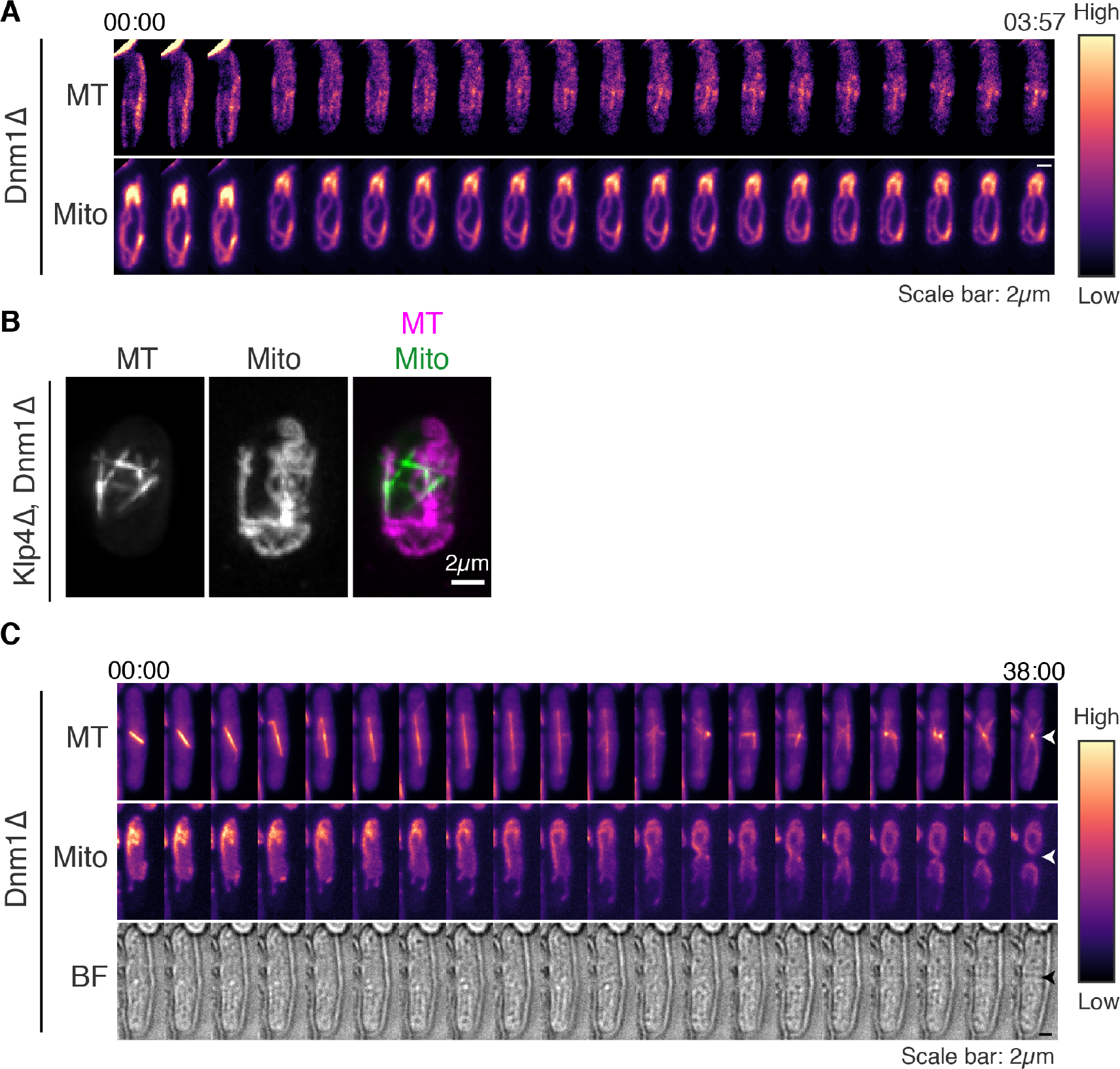
Mitochondrial fission requires Dnm1. **(A)**Montage of maximum intensity projected confocal Z-stack images of the microtubules (‘MT’, top) and mitochondria (‘Mito’, bottom) in a Dnm1Δ cell transformed with plasmid FYP2026 (see Table S1) treated with MBC. ‘00:00’ indicates time 2min after addition of MBC (50μg/ml). **(B)**Maximum intensity projections of deconvolved Z-stack images of the microtubules (left), mitochondria (centre) and their composite (right) of Klp4Δ, Dnm1Δ cells (strain VA084, see Table S1). **(C)**Montage of maximum intensity projected confocal Z-stack images of the microtubules (‘MT’, top), mitochondria (‘Mito’, center) and bright-field channel (‘BF’, bottom) in a Dnm1Δ cell undergoing cell division. The location of the septum is indicated with the white and black arrowheads. The intensity map is indicated to the right of the images. Time is indicated above the images in mm:ss.

**Figure S6:**
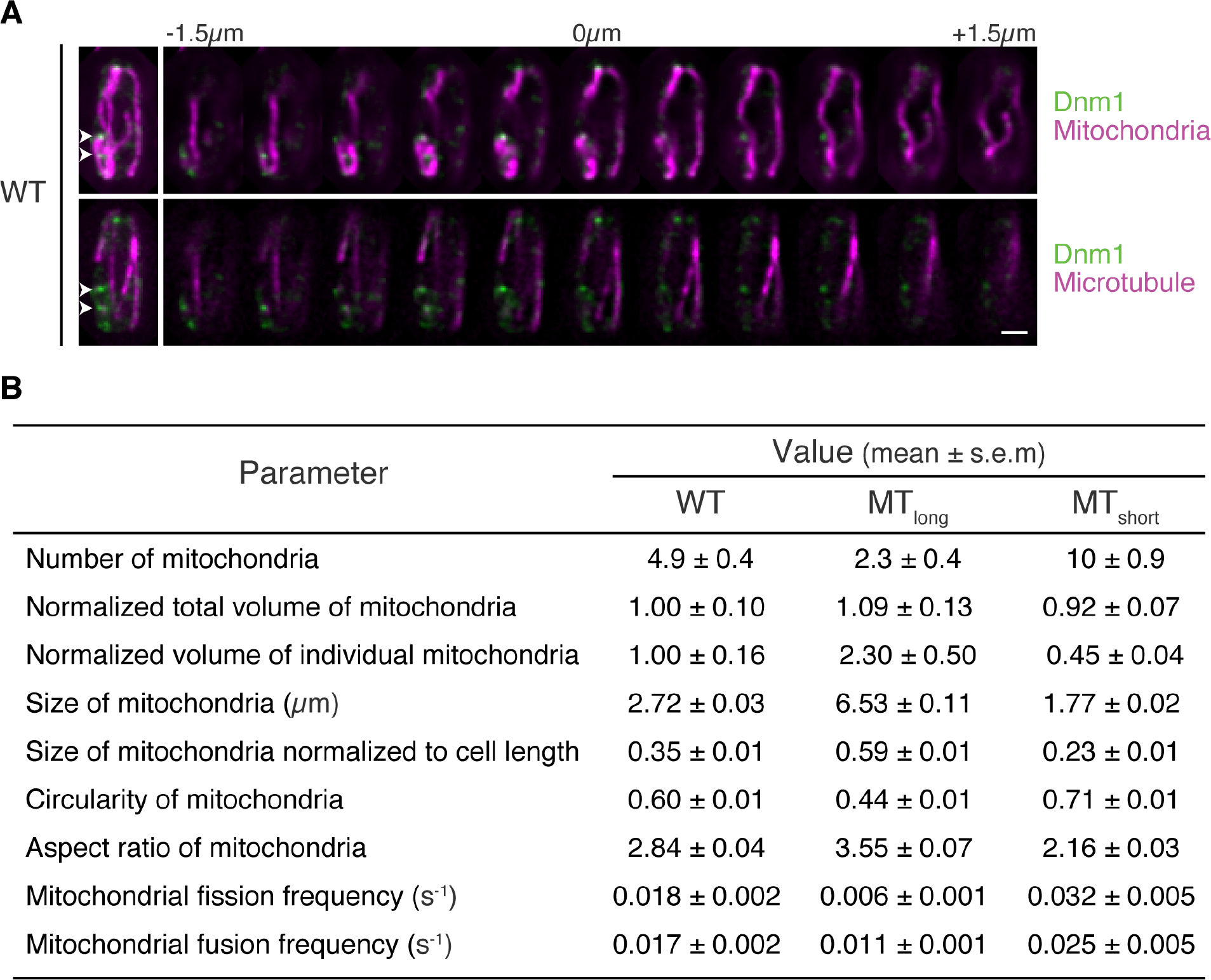
Measurement of mitochondrial dynamics. **(A)**Maximum intensity projections of deconvolved composite Z-stack images of Dnm1 and mitochondria (green and magenta respectively, top left) and Dnm1 and microtubules (green and magenta respectively, bottom left) in a wild-type cell expressing fluorescent Dnm1 and tubulin (strain MTY271 transformed with pREP41-Dnm1-Cterm-GFP, see Table S1), and stained with Mitotracker Deep Red. A majority of Dnm1 spots did not localize to mitochondria bound to the microtubule, but localize on the mitochondria away from the microtubule (white arrowheads). A montage of each 0.3μm z-slice is represented to the right of the maximum intensity projected images. The distance from the central slice (‘0μm’) is indicated above the montage. Scale bar represents 2μm. **(B)**Measured parameters relating to mitochondrial numbers and morphology in WT, MT_long_ and MT_short_ interphase cells represented as mean ± s.e.m.

### Supplementary files

**Movie S1.**Live-cell confocal microscopy of MBC-treated interphase fission yeast cells with the mitochondria stained with Mitotracker. Warmer colours indicate higher intensities. Images were acquired at 2min intervals. Time 00:00 (mm:ss) indicates time of addition of MBC. Imaging was resumed 15min after MBC addition (’15:00’). Scale bar represents 2μm. This movie corresponds to Fig. S1C.

**Movie S2.**3D projection of microtubules (green) and mitochondria (magenta) in a wild-type cell (strain KI001, see Table S1). This movie corresponds to Fig. 1A.

**Movie S3.**3D projection of microtubules (green) and mitochondria (magenta, stained with Mitotracker) in a Klp5Δ/Klp6Δ cell (strain G3B, see Table S1). This movie corresponds to Fig. 1A.

**Movie S4.**3D projection of microtubules (green) and mitochondria (magenta, stained with Mitotracker) in a Klp4Δ cell (strain G5B, see Table S1). This movie corresponds to Fig. 1A.

**Movie S5.**Mitochondrial dynamics in a wild-type cell (strain KI001, see Table S1) stained with Mitotracker. A 5-slice Z-stack with step size of 0.5μm was obtained every 12s. The movie represents the maximum-intensity projected images of the stacks obtained. Warmer colours indicate higher intensity. The closed white arrowhead points to a fusion event and the open white arrowhead points to a fission event. Scale bar represents 2μm. This movie corresponds to Fig. 2A.

**Movie S6.**Mitochondrial dynamics in a Klp5Δ/Klp6Δ cell (strain G3B, see Table S1) stained with Mitotracker. A 5-slice Z-stack with step size of 0.5μm was obtained every 12s. The movie represents the maximum-intensity projected images of the stacks obtained. Warmer colours indicate higher intensity. The closed white arrowhead points to a fusion event. Scale bar represents 2μm. This movie corresponds to Fig. 2A.

**Movie S7.**Mitochondrial dynamics in a Klp4Δ cell (strain G5B, see Table S1) stained with Mitotracker. A 5-slice Z-stack with step size of 0.5μm was obtained every 12s. The movie represents the maximum-intensity projected images of the stacks obtained. Warmer colours indicate higher intensity. The open white arrowhead points to a fission event. Scale bar represents 2μm. This movie corresponds to Fig. 2A.

**Movie S8.**Mitochondrial dynamics in a wild-type cell treated with MBC (strain KI001, see Table S1) stained with Mitotracker. A 5-slice Z-stack with step size of 0.5μm was obtained every 12s. The movie represents the maximum-intensity projected images of the stacks obtained. Warmer colours indicate higher intensity. The open white arrowhead points to a fission event. Scale bar represents 2μm. This movie corresponds to Fig. 3A.

**Movie S9.**Mitochondrial dynamics in a Klp5Δ/Klp6Δ cell treated with MBC (strain G3B, see Table S1) stained with Mitotracker. A 5-slice Z-stack with step size of 0.5μm was obtained every 12s. The movie represents the maximum-intensity projected images of the stacks obtained. Warmer colours indicate higher intensity. The closed white arrowhead points to a fusion event. Scale bar represents 2μm. This movie corresponds to Fig. 3A.

**Movie S10.**3D projection of microtubules (green) and mitochondria (magenta) in a wild-type mitotic cell (strain KI001, see Table S1). This movie corresponds to Fig. 4A.

**Movie S11.**Microtubule dynamics (left) and mitochondrial dynamics (right) in a wild-type cell (KI001, see Table S1) at the onset of mitosis. A 5-slice Z-stack with step size of 0.5μm was obtained every 2min. The Movie represents the maximum-intensity projected images of the stacks obtained. Warmer colours indicate higher intensity. Scale bar represents 2μm. The open white arrowheads point to fission events. This movie corresponds to Fig. 4B.

**Movie S12.**3D projection of microtubules (green) and mitochondria (magenta) in an Mmb1 Δ cell (strain PT2244, see Table S1). This movie corresponds to Fig. 6A.

**Movie S13.**3D projection of microtubules (green) and mitochondria (magenta) in an Klp5Δ/Klp6Δ -Mmb1 Δ cell (strain VA076, see Table S1). This movie corresponds to Fig. 6A.

